## Supplementary Figures for "Two opposing roles for Bmp signalling in the development of electrosensory lateral line organs"

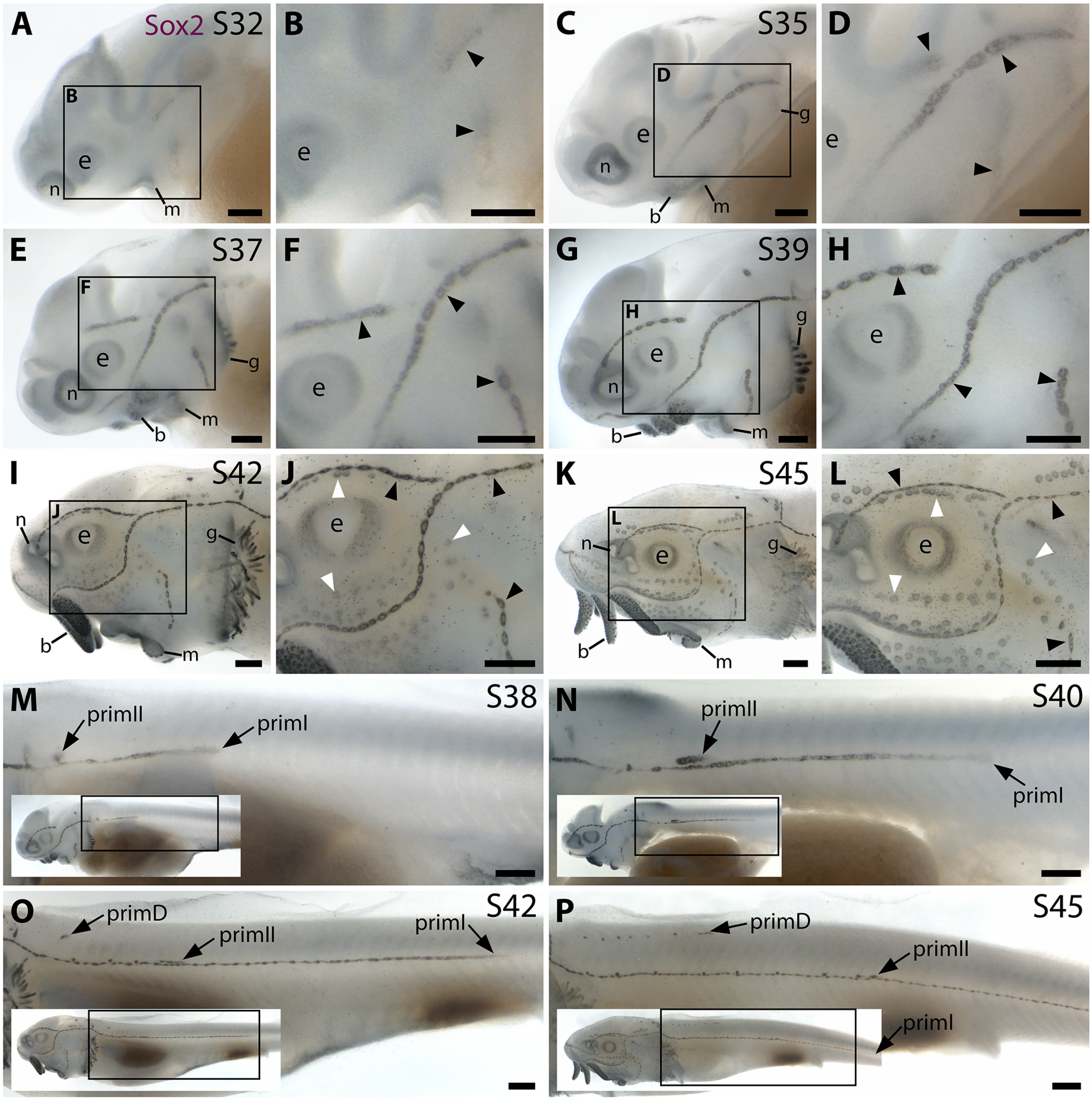


**Supplementary Figure S1. Sox2 expression shows the time-course of sterlet lateral line organ development.** Sterlet embryos and yolk-sac larvae immunostained at selected stages for Sox2, which labels lateral line primordia/sensory ridges and supporting cells in lateral line organs. Black arrowheads indicate examples of developing neuromasts; white arrowheads indicate examples of developing ampullary organs. Non-lateral line expression is seen around the mouth and nares, in the eye, gill filaments and taste buds on the barbels. Scattered individual Sox2-positive skin cells are most likely Merkel cells. (**A-D**) At stage 32 (A,B) and stage 35 (C,D), Sox2 expression is seen in the otic/anterodorsal and anteroventral lateral line primordia, with a ring pattern around developing neuromast primordia. (**E-H**) At stage 37 (E,F) and stage 39 (G,H), Sox2 is expressed at the edges of neuromasts and in interneuromast cells. (**I-L**) At stage 42 (I,J) and stage 45 (K,L), Sox2 expression is expressed at the periphery of neuromasts and ampullary organs, with stronger expression in neuromasts. The images shown in K,L were previously published in Minařík et al. (2024a). (**M-P**) Sox2 expression on the trunk at stage 38 (M), stage 39 (N), stage 42 (O) and stage 45 (P) shows the progression of the migrating lateral line primordia (primI, primII and primD). Low-power insets show the location of these trunk regions. Sox2 is expressed only weakly in primI (e.g., M,N), but strongly in primI-deposited neuromasts and interneuromast cells (M-P). PrimII is located a little dorsal to the primI-deposited line (M-P), and primII-deposited neuromasts are slightly offset dorsally from the line of primI-deposited neuromasts (O,P). Sox2 expression only reveals neuromasts deposited by primII and primD, not interneuromast cells (O,P; compare with the primI-deposited line in M-P). Abbreviations: b, barbel; e, eye; g, gill filaments; m, mouth; n, naris; prim, migrating lateral line primordium (primI, primary; primII, secondary; primD, dorsal); S, stage. Scale bar: 250 μm.

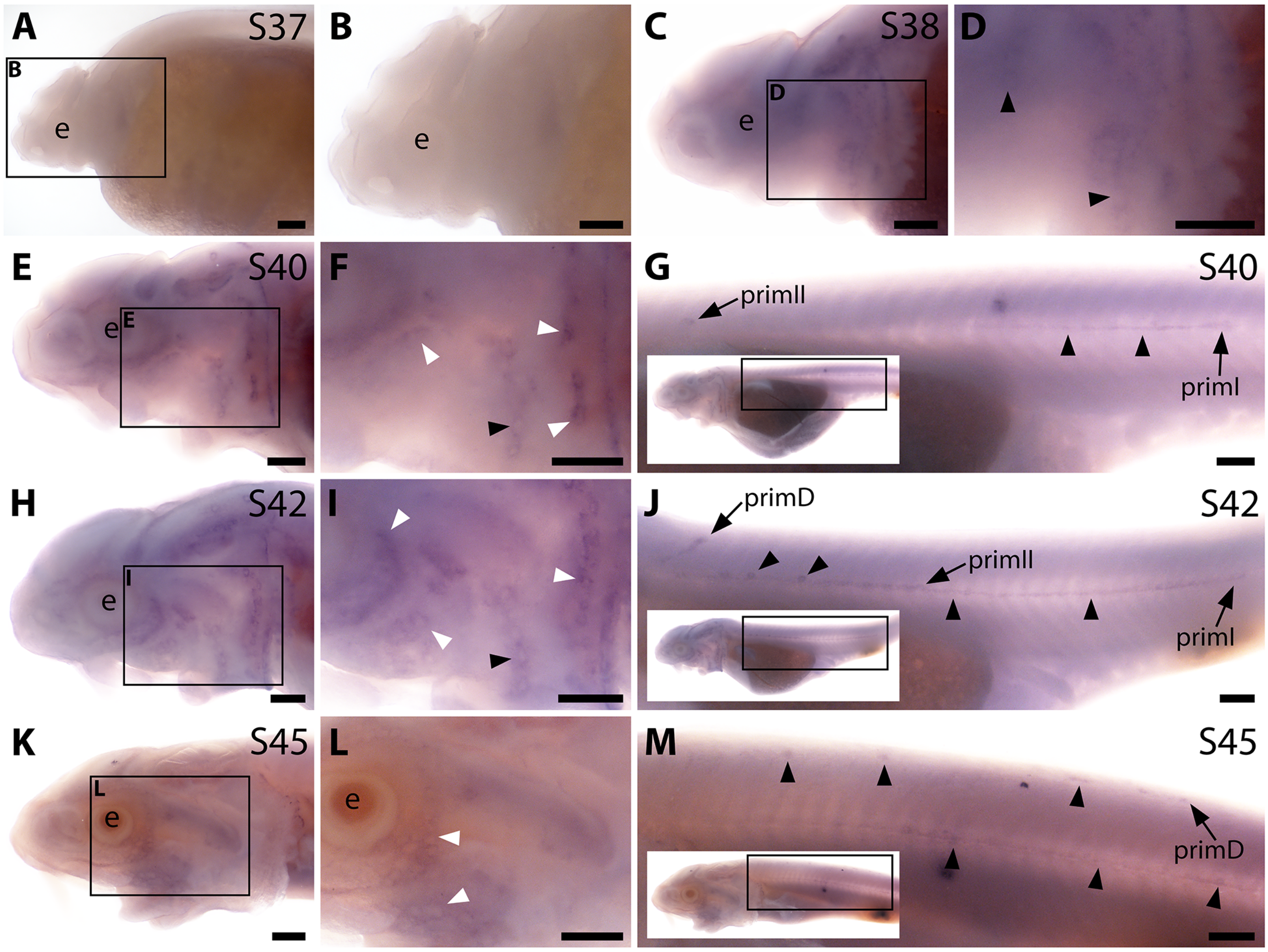


**Supplementary Figure S2. Sterlet *Acvr2a* is expressed in developing neuromasts and ampullary organs.** *In situ* hybridisation in sterlet for the type II receptor gene *Avcr2a*. Black arrowheads indicate examples of developing neuromasts; white arrowheads indicate examples of developing ampullary organs. For trunk images, boxes on low-power insets delineate the regions shown. (**A,B**) At stage 37, there is no detectable *Avcr2a* expression. (**C,D**) At stage 38, faint expression is seen in developing neuromast regions. (**E-J**) At stage 40 (E-G) and stage 42 (H-J), *Avcr2a* is expressed in developing ampullary organ primordia and neuromasts on the head (E,F) as well as in the migrating lateral line primordia and neuromasts on the trunk. (**K-M**) By stage 45, only faint *Acvr2a* expression remains around some ampullary organs on the head (K,L), although expression is still seen in the migrating primordia and neuromasts on the trunk (M). Abbreviations: e, eye; prim, migrating lateral line primordium (prim**I**, primary; prim**II**, secondary; primD, dorsal); S, stage. Scale bar: 250 μm.

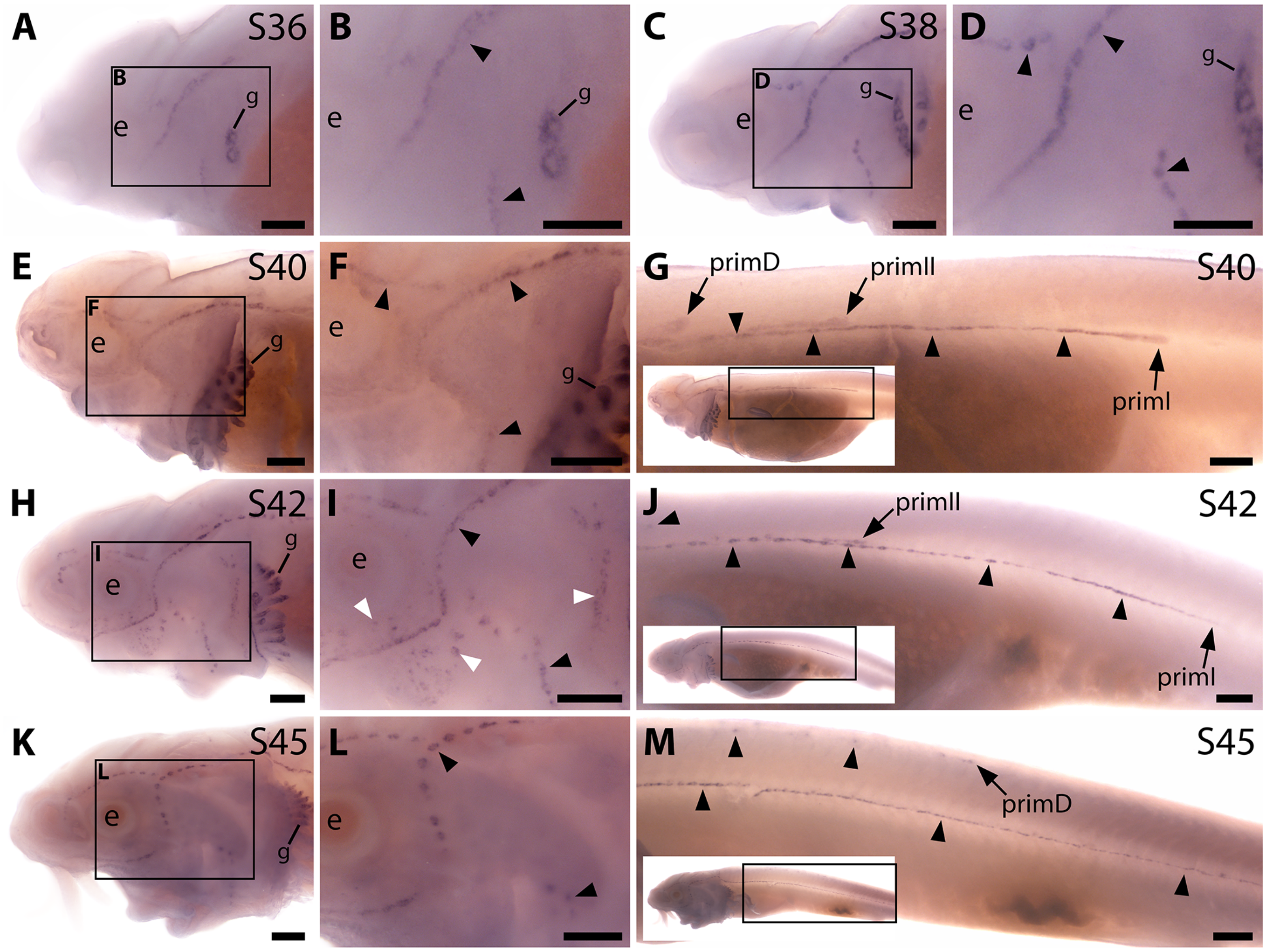


**Supplementary Figure S3. Sterlet *Sostdc1* is expressed in neuromasts but only transiently in ampullary organs.** *In situ* hybridisation in sterlet for *Sostdc1*, encoding a secreted dual Wnt/Bmp antagonist. Black arrowheads indicate examples of developing neuromasts; white arrowheads indicate examples of developing ampullary organs. For trunk images, boxes on low-power insets delineate the regions shown. Non-lateral line expression is seen in developing gill filaments. (**A-D**) *Sostdc1* expression is seen in neuromasts on the head from stage 36 (A,B), persisting at stage 38 (C,D). (**E-G**) At stage 40, expression is maintained in neuromasts on the head (E,F) and is also visible in the three migrating lateral line primordia on the trunk and in trunk neuromasts (G). (**H-J**) At stage 42, *Sostdc1* expression persists in cranial neuromasts and is now also seen in ampullary organs (H,I). On the trunk, expression persists in the migrating primordia and neuromasts (J). (**K-M**) By stage 45, only neuromast expression is observed on the head (K,L). On the trunk, expression continues in the migrating primordia and neuromasts (M). Abbreviations: e, eye; prim, migrating lateral line primordium (primI, primary; primII, secondary; primD, dorsal); S, stage. Scale bar: 250 μm.

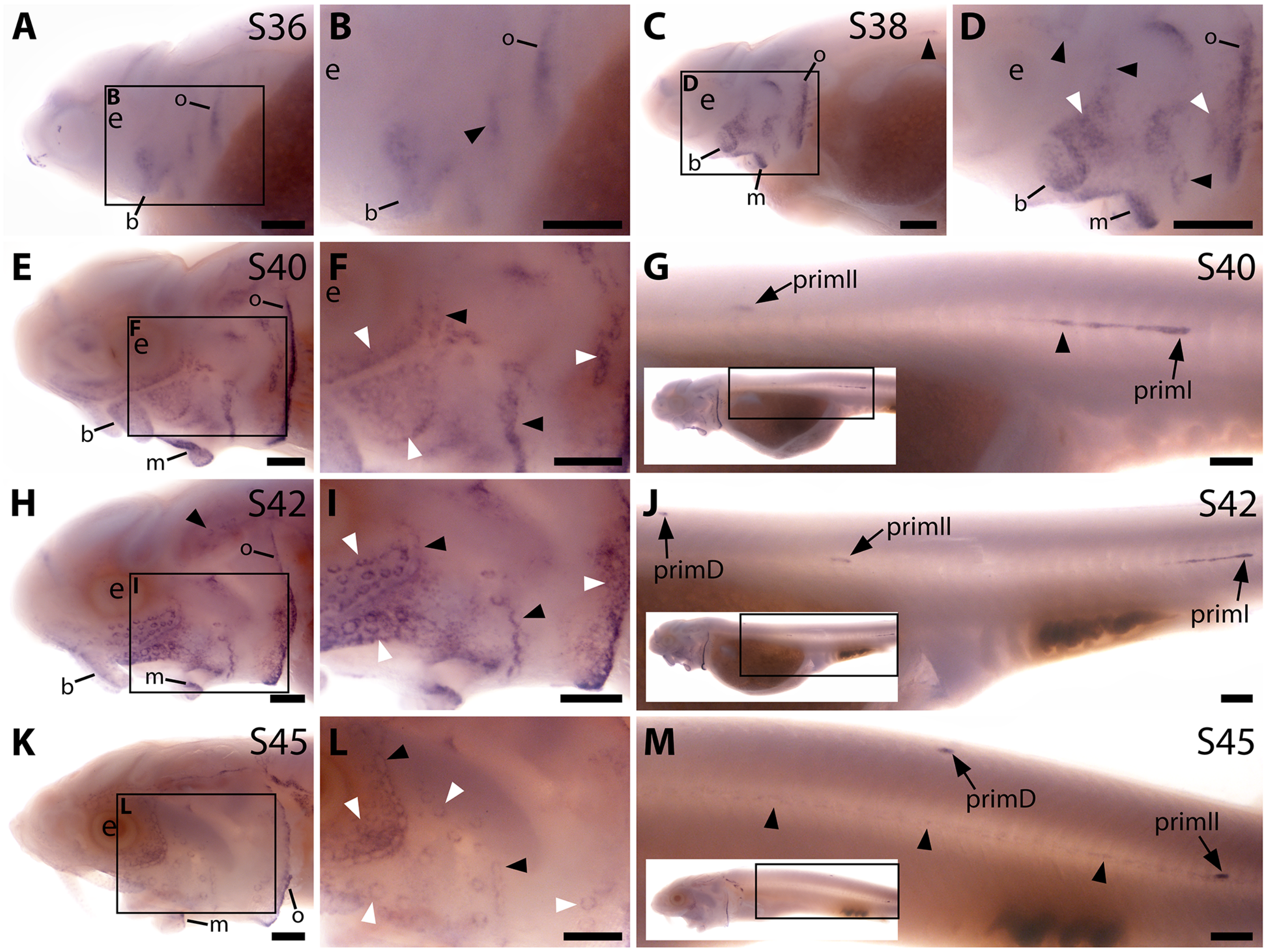


**Supplementary Figure S4. Sterlet *Apcdd1* is expressed in ampullary organs and neuromasts during development.** *In situ* hybridisation in sterlet for *Apcdd1*, encoding a secreted dual Wnt/Bmp antagonist. Black arrowheads indicate examples of developing neuromasts; white arrowheads indicate examples of developing ampullary organs. For trunk images, boxes on low-power insets delineate the regions shown. (**A,B**) At stage 36, *Apcdd1* expression is seen in the region of the preopercular neuromast line, as well as in developing barbel regions, the opercular edge and around the mouth. (**C,D**) By stage 38, expression in all these locations is more distinct. (**E-J**) At stage 40 (E-G) and stage 42 (H-J), *Apcdd1* is expressed around ampullary organ primordia and neuromasts on the head (E,F,H,I) and is also seen on the trunk in primI and primII and a fairly short line of trailing cells behind primI (G,J). (**K-M**) At stage 45, this expression pattern largely persists on the head, although appears to be fading in the ventral infraorbital ampullary organ field (K,L). On the trunk, faint expression is seen along the main body line, with strong expression in primD and primII (M). Abbreviations: b, barbels; e, eye; m, mouth; o, operculum edge; prim, migrating lateral line primordium (primI, primary; primII, secondary; primD, dorsal); S, stage. Scale bars: 250 μm.

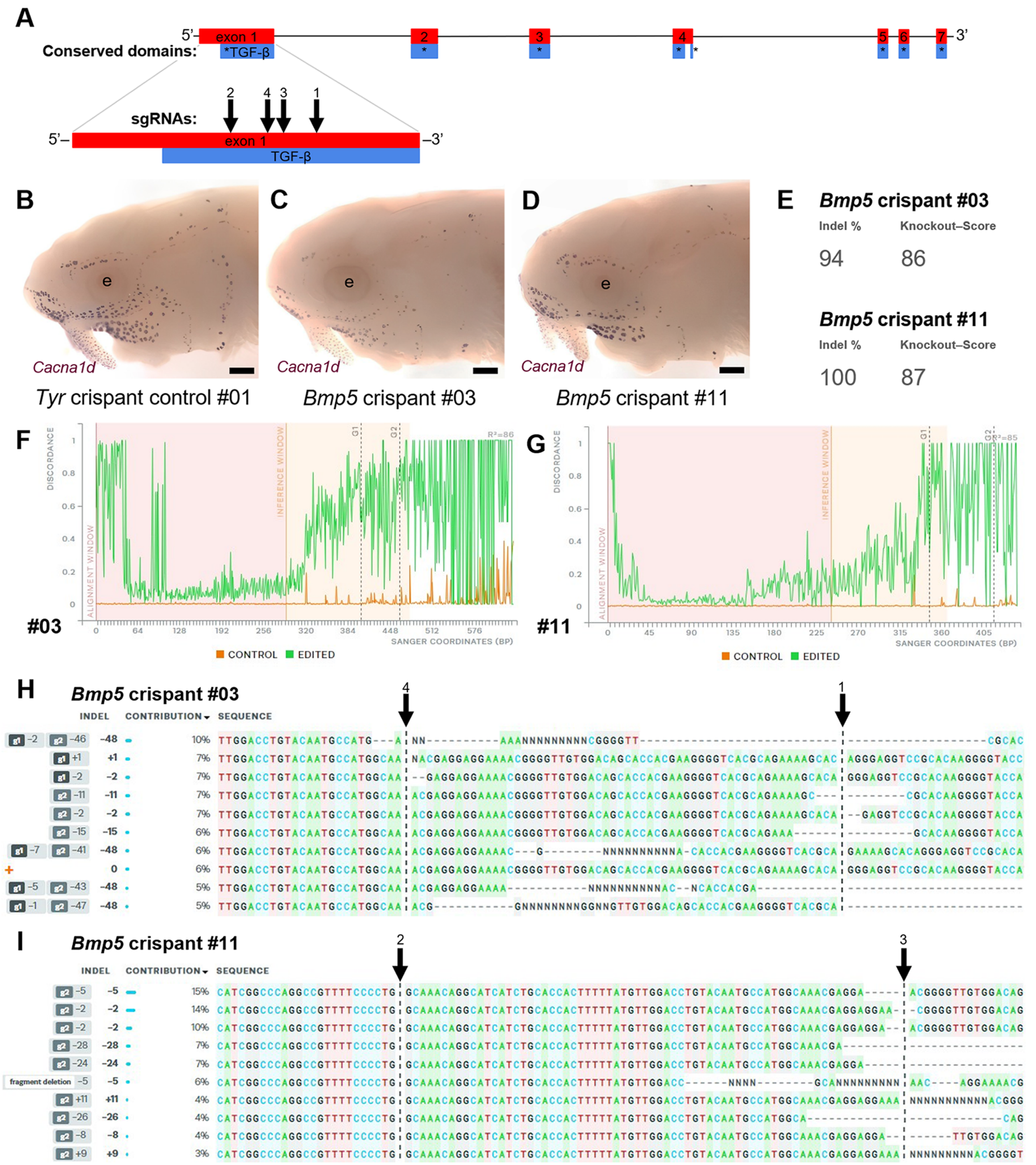


**Supplementary Figure S5. Examples of successful disruption of sterlet *Bmp5* by CRISPR/Cas9-mediated mutagenesis in G0-injected embryos.** (**A**) Schematic showing the exon structure of the sterlet *Bmp5* gene relative to conserved domains and the target sites of *Bmp5* sgRNAs (Table 1). (**B-D**) Sterlet crispants at stage 45 after *in situ* hybridisation for the hair cell and electroreceptor marker *Cacna1d* (also expressed in taste buds on the barbels). In comparison to the control *Tyr* crispant (B), the two *Bmp5* crispants (C,D) have fewer ampullary organs (this is particularly clear on the operculum). These images are also shown in Figure 4A,C,E. (**E-I**) Outputs are shown from Synthego's 'Inference of CRISPR Edits' (ICE) tool (Conant et al., 2022). This tool was used to analyse Sanger sequence data for the targeted region of genomic DNA extracted from the trunk of each of the *Bmp5* crispants shown in panels C and D. 'Indel %' (E) gives the percentage of insertions and/or deletions among the inferred sequences in the CRISPR-edited population. 'Knockout-Score' (E) shows the proportion of indels introducing a frameshift, or that are at least 21 bp in length. Discordance plots (F,G) show the level of discordance between the control sample trace file (orange) and the edited sample Sanger trace file (green). The expected cut sites for the sgRNAs are shown by vertical dotted lines. A successful CRISPR edit is indicated by the increase in discordance near the expected cut site. Panels H and I show nucleotide sequences from the Sanger trace files and their inferred relative contributions to the edited mosaic population. Vertical dotted lines indicate the expected cut sites for the sgRNAs. The wild-type sequence (0) is marked by an orange "+" symbol in H, but is absent in panel I because 100% of the sequence was edited in this crispant. Abbreviation: e, eye. Scale bar: 250 μm.
